# Odour of domestic dogs infected with *Leishmania infantum* is attractive to female but not male sand flies: evidence for parasite manipulation

**DOI:** 10.1101/2020.11.09.374207

**Authors:** Monica E. Staniek, James G.C. Hamilton

## Abstract

Globally visceral leishmaniasis (VL) causes thousands of human deaths every year. In South America, the etiologic agent, *Leishmania infantum*, is transmitted from an infected canine reservoir to human hosts by the blood-feeding activity of the sand fly vector, predominantly, *Lutzomyia longipalpis*. Previous evidence from model rodent systems have suggested that the odour of infected hosts is altered by the parasite making them more attractive to the vector leading to an increased biting rate and improved transmission prospects for the pathogen. However, the effect of *Le. infantum* infection on the attractiveness of naturally infected dogs which are integral to human infection, has not been assessed.

Hair collected from infected and uninfected dogs residing in a VL endemic city in Brazil. was entrained to collect the volatile chemical odours present in the headspace. Female and male *Lu. longipalpis* sand flies were offered a choice of infected or uninfected odour in a series of behavioural experiments. Control experiments established that female and male *Lu. longipalpis* were equally attracted to uninfected dog odour, female *Lu. longipalpis* were significantly more attracted to infected dog odour than uninfected dog odour in all 15 experimental replicates (average 45.7±0.87 females attracted to infected odour; 23.9±0.82 to uninfected odour; paired T-test, *P*=0.000). Male *Lu. longipalpis* did not significantly prefer either infected or uninfected odour (average 36.1±0.4 males to infected odour; 35.7±0.6 to uninfected odour; paired T-test, *P*=0.722). A significantly greater proportion of females chose the infected dog odour compared to the males (paired T-test, P=0.000).

The results show that dogs infected with *Le. infantum* are significantly more attractive to blood-feeding female than male sand flies. This is strong evidence for parasite manipulation of the host odour in a natural transmission system and indicates that infected dogs may have a disproportionate significance in maintaining infection in canine and human infection.

**Author Summary:** Visceral leishmaniasis (VL) is a disease caused by the Protist parasite *Leishmania infantum*. In Brazil and other South and Central American countries, the parasite is transmitted by the blood-feeding activity of infected female *Lutzomyia longipalpis* sand flies. The disease leads to thousands of human cases and deaths every year. Domestic dogs are the reservoir of infection for humans therefore understanding the effect of infection on dogs is important in developing an understanding of the epidemiology of the disease. Although previous studies on rodent models of *Le. infantum* infection have shown that infected Golden Hamsters are more attractive to *Lu. longipalpis* the attractiveness of naturally infected dogs to the insect vector has not been previously been investigated. In this study we showed that the odour of infected dogs is significantly more attractive to female sand flies which can transmit the pathogen than to male sand flies which do not. This clear-cut difference in attraction of female and males suggests that the females are preferentially attracted by parasite infected hosts and may lead to enhanced infection and transmission opportunities for the parasite.

## Introduction

In South America the protist *Leishmania (Le.) infantum* (Cunha & Chagas) (Kinetoplastida: Trypanosomatidae) causes the disease visceral leishmaniasis (VL) in humans and dogs (*Canis lupus familiaris*) [1]. The intracellular parasite, which is a pathogen of the immune system, primarily affects the reticuloendothelial system, surviving and multiplying in macrophage and dendritic cells [2]. In humans, VL is characterized by prolonged irregular fever, splenomegaly, hepatomegaly, pancytopenia and weight loss and in the absence of treatment the case fatality rate is 90% [3].

Globally, approximately 50,000 to 90,000 new cases of VL occur each year with more than 95% of them occurring in ten countries: Brazil, China, Ethiopia, India, Iraq, Kenya, Nepal, Somalia, South Sudan and Sudan [4]. In the Americas, leishmaniasis is a significant public health problem due to its’ morbidity, mortality and broad geographical distribution [5]. From 2001 to 2015, 52,176 cases of visceral leishmaniasis were registered in the Americas, with 96.4% of these cases (50,268) reported from Brazil [5, 6]. Overall, the burden of the disease (age-standardized DALY) in Brazil has almost doubled during the period from 1990 to 2016 [7].

In Brazil *Leishmania infantum* infection is a zoonosis, with canines as the primary reservoir hosts. Humans, which are considered to be a dead-end host for the parasite [8], become infected when they are fed on predominantly by infected female sand flies of the *Lutzomyia longipalpis s.l.* (Lutz & Neiva) (Diptera: Psychodidae) species complex [9]. In dogs, VL is characterised by dermatitis, lymphadenomegaly, general muscular atrophy, and renal disease [10].

Host seeking haematophagous insects orientate towards and identify potential host animals by combinations of visual, thermal tactile and chemical (host odour) cues produced by the host [11]. Together these stimuli, along with the individual response of female insects, dictate the likelihood that a haematophagous insect will successfully obtain a blood meal [11–13]. For temporary ectoparasites (e.g. mosquitoes, triatomines, midges and sand flies) host odour is of primary importance in host location. Parasite manipulation of the host can enhance transmission opportunities in favour of the parasite. There is now significant evidence that parasite infection of the host animal alters both the vector host choice behaviour as well as vector biting rate and blood-meal size [14]. The choice to bite an infected or uninfected host is key for pathogen transmission and numerous empirical studies show that pathogens can modify the scent of infected hosts to attract vectors [15].

Mosquito vectors are more attracted to hosts, including humans, mice and birds, infected with malaria than to uninfected hosts [16–20]. Humans infected with *Plasmodium falciparum* are more attractive to *Anopheles gambiae s.l.* than uninfected humans. The increased attraction is most pronounced when the parasite is in the most transmissible (gametocyte) stage [21] and the increased host parasitaemia may partially, at least, be induced by mosquito biting activity which leads in turn to higher transmission rates [22]. The difference in attractiveness has been related to the presence of a parasite produced isoprenoid precursor, (*E*)-4-hydroxy-3-methyl-but-2-enyl pyrophosphate (HMBPP) in host blood which has been shown to trigger an enhanced release of attractants which in turn increase the likelihood of an infected human host being bitten by the vector [23].

In contrast only a limited number of studies have been carried out to determine if *Leishmania* parasites manipulate host odour. In olfactometer choice experiments volatile odours collected from hamsters infected with *Le. infantum* (MHOM/BR/74/PP75) were found to be more attractive to female *Lu. longipalpis* sand flies than uninfected hamsters [24]. A subsequent laboratory study investigated the change in attractiveness of individual hamsters infected with *Leishmania infantum* and showed that the odour of a significant number of hamsters became significantly more attractive to the vector after the *Leishmania* parasites had reached the metacyclic infection stage compared to the initial amastigote infection stage [25]. A coupled gas chromatography-mass spectrometry (GC/MS) and statistical analysis (principal component analysis, PCA) compared the volatile organic chemical profile obtained from the headspace of the hair of a small group of Brazilian dogs infected with *Le. infantum* with an uninfected group of dogs and found a significant difference in the odour profiles of the two groups [26, 27]. The key discriminatory volatile components identified by the PCA were characterised by conventional analytical methods [27] and in a behavioural study were found to be attractive albeit at very high concentrations (≥50% purity) predominantly to male *Lu. longipalpis* [28].

None of these previous studies conclusively demonstrated that the parasite *Le. infantum* enhanced the attractiveness of the odour of the natural canine reservoir host to the sand fly vector; the experiments with hamsters were carried out with a non-natural host/parasite system and the experiments with canine odour, although showing a difference in odour profile between infected and uninfected dogs, did not test if the infected dog odour was more or less attractive than the uninfected dog odour.

We therefore carried out a study on the odours collected from the hair of dogs in an olfactometer, to determine if female or male *Lu. longipalpis* had a preference for either infected or uninfected odour.

## Results

### Female sand fly response to infected and uninfected dog odour

In all 15 individual experimental replicates, female *Lu. longipalpis* were significantly attracted to the infected dog odour compared to the uninfected odour (Fig 1A; S1 Table).

**Fig 1.**
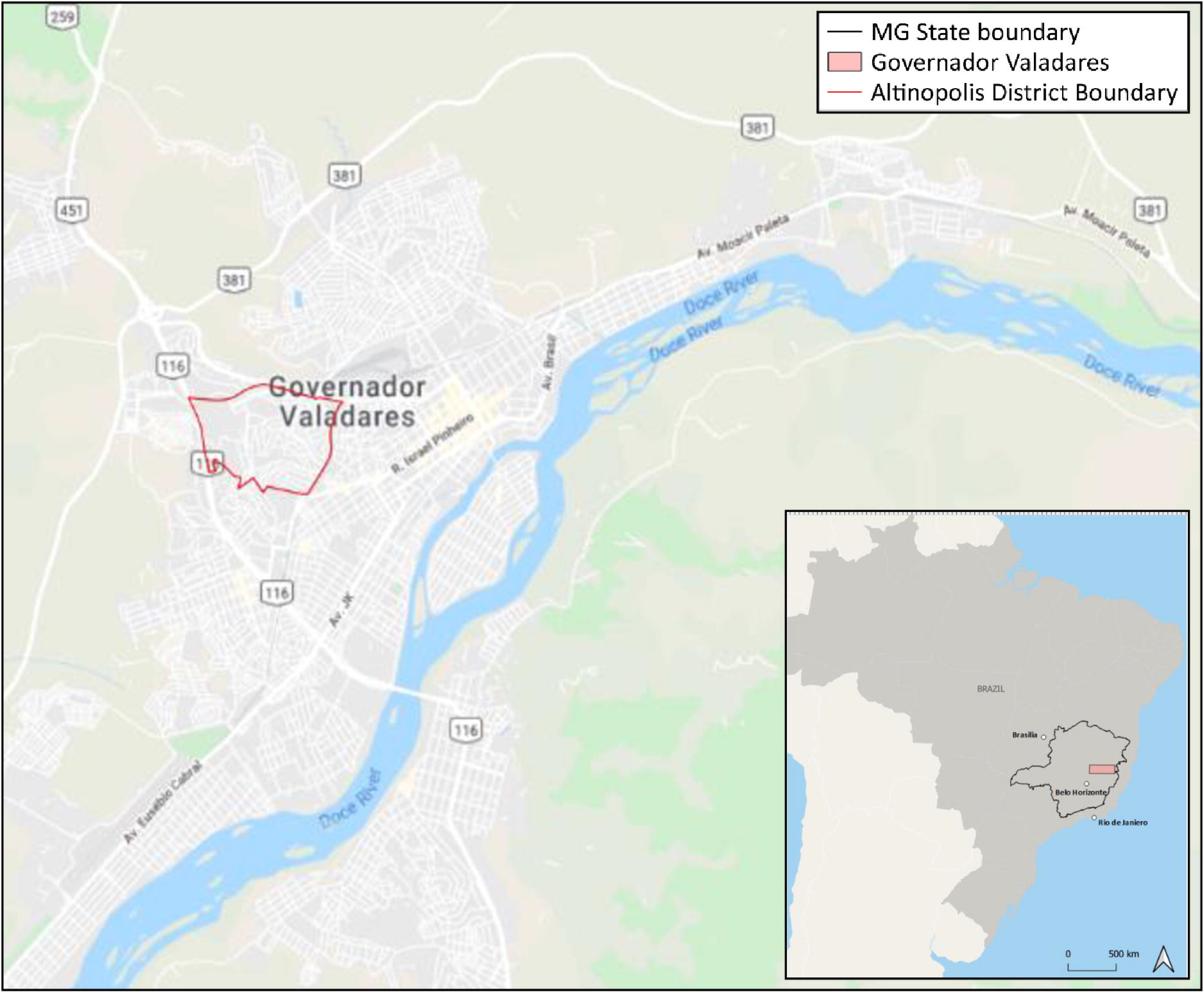
Numbers of female (A) and male (B) *Lutzomyia longipalpis* attracted to volatiles collected from infected and uninfected dogs. Numbers of female (A) or male (B) *Lu. longipalpis* attracted to either the infected (grey bars) or uninfected (hatched bars) dog odour presented in the Y-tube choice olfactometer. The numbers attracted to each side are given beside each bar. The dog odour pairs are indicated on the left axis, the first dog in each pair is the infected animal. The superscripts indicate if the dog infection status was known to the experimenter or not; α for the female sand fly bioassay indicates that the status was known β indicates it was not; similarly, γ for the male sand fly assays indicates that it was known and δ indicates it was not. Statistical significance was analysed by binomial test comparing the actual number of sand flies that responded to infected odour compared to the number that responded to the uninfected odours (\**P*≤0.05, \*\**P*≤0.01, \*\*\**P*≤0.001, \*\*\*\**P*≤0.0001, ns=not significant).

**Fig. 1.**
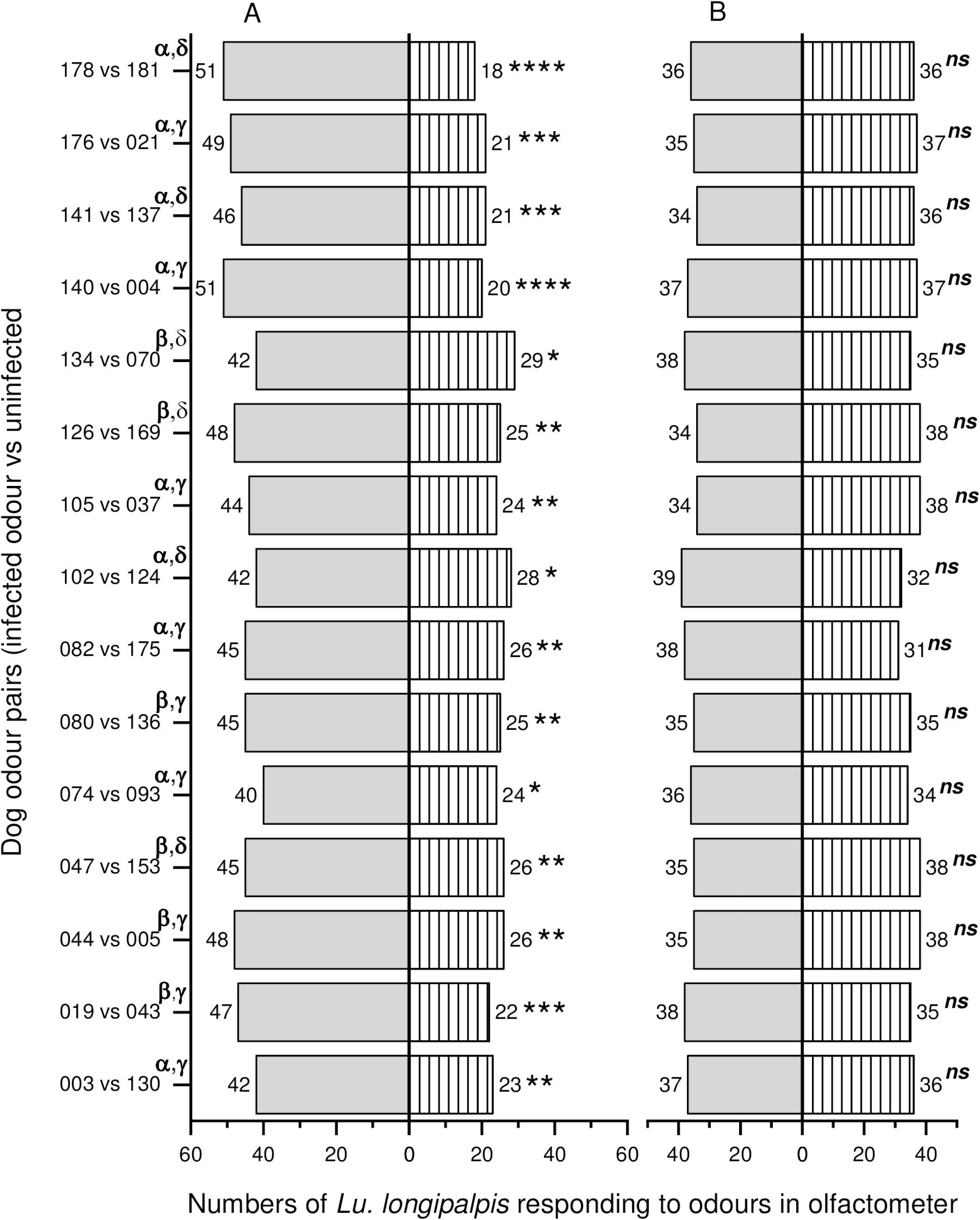
The municipality of Governador Valadares (Minas Gerais State Brazil) and its location in Brazil. The hair samples were from dogs recruited in the Altinopolis district of the city, bordered with a red line. Map in was constructed in the QGIS 3.10, using OpenStreetMap (Map data copyrighted OpenStreetMap contributors and available from https://www.openstreetmap.org) as base map.

We combined the data from the blinded and unblinded experiments for each of the infected and uninfected data sets as there was no significant difference between them [infected dog odour; unblinded 45.6±1.3 vs blinded 45.83±1.0 (2-sample T-test, *P*=0.869); uninfected dog odour; unblinded 22.8±1.1vs blinded 25.5±1.0 (2-sample T-test; *P*=0.0874)] and then compared the response to each (mean response to infected odour 45.7±0.87; mean response to uninfected odour 23.9±0.82). Female sand flies were significantly attracted to the infected odour (paired T-test, *P*=0.000).

The data were normally distributed (Anderson-Darling *P*>0.1 and Kolmogorov-Smirnov P>0.1).

### Male sand fly response to infected and uninfected dog odour

In all 15 of the individual experimental replicates, male *Lu. longipalpis* were not significantly more attracted to the infected dog odour entrainment extracts compared to the uninfected dog odour (Fig 1B; S2 Table).

We combined the data from the blinded and unblinded experiments for each of the infected and uninfected data sets as there was no significant difference between them [infected dog odour; unblinded 36.1±0.5 vs blinded 36.0±0.9 (2-sample T-test, *P*=0.913); uninfected dog odour; unblinded 35.8±0.8 vs blinded 35.8±0.9 (2-sample T-test; *P*=0.890)] and then compared response to each (mean response to infected odour 36.1±0.4; mean response to uninfected odour 35.7±0.6). Male sand flies were equally attracted to the infected and uninfected odours (paired T-test, *P*=0.722).

The data were normally distributed (Anderson-Darling *P*>0.1 and Kolmogorov-Smirnov P>0.1).

#### Comparison of female and male response to infected dog odour

On average significantly more of the females that responded (65.7±1.1%) were attracted to the infected dog odour than the males 50.3±0.6% of the males (T-test; *P*<0.000).

### Relationship between parasite load, discriminate value and the female and male sand fly response

There was no significant relationship between the parasite load and the proportion of females (or males) that were attracted to the infected dog hair odour [Regression analysis: females R^2^(adjusted) = 0.00%; males R^2^(adjusted) = 0.00%]. There was a slight but non-significant relationship between the previously identified discriminate values [29] and the proportion of females (or males) that were attracted to the infected dog hair odour [Regression analysis: females: R^2^(adjusted = 8.1%); males R^2^(adjusted) = 16.6%, *P*=0.074]. A correlation analysis suggested a negative relationship between the discrimination value of infected dog odour and male response (Pearson r = −0.25). Removal of the parasite load value of 853.44 parasites ml^−1^, which is a potential outlier data point, had little effect on R^2^(adjusted) for either the female or male response to parasite load however the relationship between discrimination factor became slightly more important for the female sand fly response [R^2^(adjusted) = 8.9% P=0.158] and for the males [R^2^(adjusted) = 9.4% P=0.152]. Correlation analysis suggested a moderate relationship between the discriminate value and female response (Pearson r = 0.4) and male response (Pearson r = −0.4).

### Control

Both female and male sand flies were significantly attracted to dog odour. A significantly greater proportion of female sand flies (66.5 %) were attracted to the uninfected dog volatiles compared to hexane (Binomial test; P=0.000) and of the male sand flies that responded, 62.9% were attracted to uninfected dog odour compared to the hexane control (Binomial test; P=0.000) (Table 1).

**Table 1.** Response of female and male *Lutzomyia longipalpis* to odour of uninfected dogs and hexane only in Y-tube olfactometer choice experiments.

| female sand fly response |  |  |  |  |  |
| --- | --- | --- | --- | --- | --- |
| expt | dog i.d. | uninfected dog odour | hexane | no-response | *P* |
| 1 | 138 | 48 | 21 | 11 | *** |
| 2 | 128 | 45 | 29 | 6 | * |
| 3 | 027 | 46 | 20 | 14 | *** |
|  | cumulative | 139 | 70 | 31 | **** |
| male sand fly response |  |  |  |  |  |
| expt | dog i.d. | uninfected dog odour | hexane | no-response | *P* |
| 4 | 138 | 43 | 27 | 10 | * |
| 5 | 128 | 47 | 22 | 11 | *** |
| 6 | 027 | 42 | 29 | 9 | * |
|  | cumulative | 132 | 78 | 30 | **** |

Female/male sand fly response = control experiments measuring either the response of the female or male sand flies in the Y-tube olfactometer. expt = experimental replicate; dog i.d. = dog identification number; uninfected dog odour = number of sand flies attracted to the dog odour sample; hexane = number of sand flies attracted to hexane only; no-response = number of sand flies that did not respond. Statistical significance was analysed by binomial test
comparison of number of sand flies responding to the dog odour vs hexane (\* $P \leq 0.05$ , \*\*\* $P \leq 0.001$ , \*\*\*\* $P \leq 0.0001$ ).

## Discussion

Domestic dogs, *Canis lupus familiaris*, are the primary reservoir host of the Protist parasite *Le. infantum* which is transmitted by the blood-feeding activity of female *Lu. longipalpis* sand flies in Brazil. This study has shown that the odour of dogs naturally infected with *Le. infantum* was significantly more attractive to female *Lu. longipalpis* than uninfected dog odour. In addition, the odour of infected and uninfected dogs was equally attractive to male *Lu. longipalpis*.

In each of the 15 experimental replicates where female *Lu. longipalpis* were offered a choice of infected dog odour or uninfected dog odour, the females chose the infected dog odour. Overall, 685 (mean±sem: 45.7±0.9) female sand flies chose the infected odour compared to 358 (23.9±0.8) that chose the uninfected dog odour. By comparison, in all 15 experimental replicates the male *Lu. longipalpis* did not differentiate between infected and uninfected dog odour. Overall, 541 (36.1±0.4) males chose the infected odour and 536 (35.7±0.6) chose the uninfected odour. In both sets of experiments female and male *Lu. longipalpis* were highly attracted to dog odour with only 13 and 10% respectively failing to respond to the dog odour in the olfactometer.

Previous studies using model infection systems have indicated that the odour of host animals infected with *Le. infantum* is more attractive to the sand fly *Lu. longipalpis*. A study with Golden Hamsters showed that the odour of hamsters infected with *Le. infantum* was more attractive than odour from uninfected hamsters [24] and a subsequent study showed that the odour of 6 out of 13 Golden hamsters infected with *Le. infantum* became significantly more attractive to the majority of *Lu. longipalpis* females after a prolonged period of infection [25]. Both of these studies used Golden Hamsters which would not be infected with *Le. infantum* or exposed to blood-feeding by *Lu. longipalpis* in the wild. In that study *Le. infantum* infected Golden Hamsters were not attractive to males. This is the first occasion, that we are aware of, when *Lu. longipalpis* females and males have been offered the choice of entrained headspace volatile odours from infected and uninfected dogs. The results are significant because in South and Central American, and Mediterranean countries, dogs are the reservoir of infection and are essential to the maintenance of the zoonotic transmission cycle that leads to human infection [30]. The results of this study differ in 2 important ways from those obtained from previous work using infected Golden Hamster odour [25]. First, only 6 out of 13 of the infected hamster’s odours were attractive to females after infection whereas in this study all the randomly chosen infected dog odours were attractive and second, only approximately 50% of the male *Lu. longipalpis* responded to the infected or uninfected hamster odour whereas in this study 90% of males were responsive. These results highlight the potential problems of using model rather than real systems to explore pathogenic parasite host vector interactions.

In a series of analytical chemistry studies, the odour of dogs infected with *Le. infantum* was shown to have a different odour profile to uninfected dogs [27]. Subsequently, 4 compounds identified as potential biomarkers of infection were tested in an olfactometer at either 50% or 100% concentration. In that study, the males were found to be more responsive than females in a wind tunnel olfactometer [28]. The response of both male and female *Lu. longipalpis* was also substantially lower than the response seen in this study. It is not possible to be sure if the observed differences are because of differences in olfactometer design, small sample size or are related to the range of different compounds and concentrations presented to the sand flies and which differed substantially between the two studies.

Although both male and female *Lu. longipalpis* were attracted to uninfected dog odour only the female sand flies showed an enhanced attraction to infected dog odour. The differences between the female and male sand fly response to infected dog odour observed in this study may also be important to transmission of the parasite. *Lutzomyia longipalpis* mating behaviour is believed to be driven by male lekking behaviour in which male *Lu. longipalpis* first locate the host and establish mating aggregations on or near potential host animals [31]. Males compete with each other for territory, produce a sex/aggregation pheromone and then females are attracted by a combination of the host odour and pheromone [32–34]. The females arrive at the lekking site, choose a mate, take a blood-meal and depart [35]. In this model, males first select the host animal and the females arrive later [32, 33, 36]. The current study suggests a more nuanced situation where females can choose host animals independently of their need for associated male pheromone and it is this behaviour that can be manipulated by the parasite. The majority of sand fly species do not lek and therefore it might be possible for female *Lu. longipalpis* to also locate a host and take a blood meal without being attracted to a lek site. By not attracting male *Lu. longipalpis* to infected dogs, male aggregations would be more likely to take place on uninfected dogs (as they would be the majority of the canine population). Thus, infected female sand flies might be more likely to feed on uninfected dogs with established leks and thereby satisfying the proposed need [14] of infected vectors to feed on uninfected hosts to maintain disease transmission. To test this hypothesis in the future it will be important to determine the relative attractiveness of sex pheromone plus uninfected dog odour versus infected dog odour alone, as well as the attractiveness of infected and uninfected dog odours to infected female sand flies.

Our results showed that the parasite load in the infected dogs was not significantly related to attractiveness for female *Lu. longipalpis*. It is likely that this may be because our analysis of parasite load did not differentiate between the stage of the parasite development and thus infectivity of the animal [37]. Both these factors have been shown in other studies to be important factors in the manipulation of host attractiveness, as parasite manipulation would be most pronounced when the parasite was at a stage suitable for infection of the vector [8, 38, 39].

There was a slight positive correlation between odours that were more attractive to females and the eNose discrimination response (obtained from a previous study [29]) and a slightly greater negative correlation in the male response. This suggests that the sand flies and eNose detectors are responding to similar cues.

In the future it will be important to determine if this observed phenomenon also occurs with live infected/infectious dogs and wild *Lu. longipalpis* in the peridomestic environment where a number of other factors (e.g. odour of competing alternative animal hosts, the presence of sex/aggregation pheromone and plant odours) might contribute to, divert or dilute the response of the female sand flies to infected dog odour. In addition, synthetic sex/aggregation pheromone, formulated in controlled release lures [40] is a strong attractant that maintains the aggregations of both male and female *Lu. longipalpis* [41] while also drawing them away from chicken sheds (Gonzalez et al., 2020 In Press, Gonçalves et al., 2020 Submitted), it will be important to determine if the synthetic pheromone lure can disrupt the attractiveness of infected dogs.

It has been suggested that human odour-baited traps combined with insecticides could improve capture and kill of mosquito vectors [42–44] or cattle or pig odour could be useful for control of teste flies [45]. Our work suggests that infected dog odour may also provide opportunities to develop new vector control methods. The observed sex specific increase in attraction to infected dogs demonstrates a potential new approach for the development of a novel tool for use in vector control. Further studies are required to evaluate and implement dog odour baited traps for sand fly control [46]. The use of infected dog odour as a lure could be a widespread methodology used throughout Brazil. Combining such traps with current and proposed insecticide-based methodologies could lead to a significant reduction in the number of female sandflies [40, 47].

## Materials and Methods

### Field study site, dog recruitment and sample collection

The study was carried out on the entrained odour of hair collected from infected and uninfected dogs recruited in April 2018 in Governador Valadares (GV), a municipality of approximately 280,000 people in Minas Gerais State, Brazil (18°51’12”S W; 41°56’42”W, altitude 170m, Fig. 1) 320 km northeast of Belo Horizonte, the state capital.

GV is situated in the Rio Doce basin within the Atlantic Forest region and local topography consists of valleys and hills. The climate is Aw (tropical sub-warm and sub-dry) according to the Köppen–Geiger classification [48]. The city has an average temperature of 24.2°C (annual range 15.2 - 33°C) and the average annual rainfall of 1109 mm is concentrated between October and March [49].

The area is a focus of intense VL transmission and is also endemic for cutaneous leishmaniasis where the sand fly vector, *Lu. longipalpis* is abundant [50, 51]. In GV, 212 cases of human VL were reported between 2008-2015, the cumulative VL incidence was 7 cases per 100,000 [52] with case fatality rate estimates of 8.9%-16.3% [53, 54]. In 2017, just before this study, case fatality rate was 6.4 per 100,000 [55]. Canine infection reported for the 2008-2012 period was 29% (8,622/29,724) in GV [56].

Recruitment and sampling of 133 dogs was carried out in the Altinópolos district of Governador Valadares which was chosen because of the high prevalence of canine VL (cVL) (average incidence reported in 2013 was 33.8%) [50] and the presence of a large population of household-owned dogs (n = approx. 2000) (Centro de Controle de Zoonoses (CCZ) survey). The recruitment of domestic dogs and collection of individual hair and blood samples was undertaken with the informed consent of the dog’s owners [29]. Briefly, dogs were chosen at random by walking through the area and visually identifying potential recruits and then seeking owner consent. Inclusion criteria were; dogs aged ≥3 months, without previous clinical assessment or laboratory diagnosis for cVL. Exclusion criteria were; pregnant/lactating bitches, aggressive or stray dogs. Canine dorsal hair and blood samples were obtained as previously described [29].

### Diagnosis of dogs and parasite load calculation

The diagnosis of each dog was performed previously as reported [29]. Briefly, DNA was extracted from the buffy coat of each individual dog and a PCR was performed in triplicate using the primer pair Mary F and Mary R [57]. Diagnosis was based on the outcomes of three PCR analyses. Parasite load was calculated by qPCR also using the primer pair Mary F and Mary R and a standard curve was established using extracted *Le. infantum* DNA; 1:10 serial dilutions, ranging from 0.01 to 10,000 parasites per ml and used to quantify the number of parasites in the dog blood samples [29].

### Entrainment of volatile organic compounds

Volatile organic chemicals (VOCs) from all the dog hair samples were entrained as described previously [58]. Briefly, charcoal-filtered air (15ml sec^−1^) was passed through the inlet port of a quick-fit Dreschel head into a 50ml round-bottom quick-fit glass flask which contained the dog hair (1gram). The air from the effluent port of the Dreschel head passed into a 3.5-inch long Orbo-402 Tenax-TA trap (Sigma-Aldrich Company Ltd, Dorset, UK) via a 10cm length of Teflon tubing. Hair samples were entrained for 2.5h and the collected VOCs were recovered by elution of the Orbo tube with *n*-hexane (SupraSolv grade; Merck, Germany) (2ml). A fresh Orbo tube was used for each entrainment dog hair sample. Each eluted sample was concentrated to a final volume (ca. 500μl) under nitrogen and then sealed in a clean glass Pasteur pipette and stored (−20°C) until further analysis.

### Bioassay

#### Sand flies

The sand flies used in the bioassay were from a colony maintained at Lancaster University that was originally established from individuals collected in Jacobina, Brazil (40°31′ W, 11°11′S). Flies were reared according to the method of Lawyer *et al* [59] and adults maintained in Nylon Barraud cages (18 x 18 x 18 cm) at 27°C (range 25-29°C), 95% RH with a 12:12 h (L:D) photoperiod. Female sand flies were fed sheep blood (TCS Biosciences Ltd, Buckingham, UK) via a chicken skin membrane before egg laying [60–62]. To obtain virgin female and male *Lu. longipalpis* for use in the bioassays newly eclosed adults were separated within 6 h of emergence (before rotation of external male genitalia) and fed on saturated sucrose solution only. They were then held in a Barraud cage, kept inside a plastic bag to maintain humidity at 27°C with access to sugar (*ad libidum*) in the insectary for 5 to 7 days until used in the bioassay.

Bioassays were performed as previously described [58]. One hour prior to the start of the experiment the sand fly holding cage was moved into the bioassay room where the plastic bag was removed and the sand flies allowed to acclimatise to the room conditions (temp; mean 27°C, range 25-29°C: humidity; mean 75%, range 68-78%). All bioassays were performed at the same time of day to minimise any potential effect of circadian rhythms on flight activity [63].

#### Y-tube olfactometer

The Y-tube olfactometer and associated Teflon tubing were as previously described [34, 58]. This apparatus allowed us to enter the entrained dog odour solutions into the air flowing through the Teflon tubing connected to the olfactometer arm. The solution was injected onto a rolled up filter paper disk (Grade 1; 20mm diam., Whatman International Ltd., Maidstone, United Kingdom) via a small hole drilled in the wall of the tubing. A flow of clean air (ZeroGrade; BOC address) through the apparatus was controlled by a two-stage cylinder regulator valve (BOC Series 8500 Air Regulator BOC Lancaster, UK) and a rotameter (Sigma-Aldrich Company Ltd, UK), adjusted to 5ml sec^−1^ and periodically confirmed using a bubble meter. All connections between Teflon tubing components were made airtight by sealing with Teflon tape (Sigma-Aldrich Company Ltd, UK). Glass wool was inserted into the end of each Y-tube olfactometer arm to prevent flies from escaping into the apparatus.

All Teflon tubing and glassware was cleaned 24h prior use. Glassware was washed with 10% Teepol solution, distilled water and then acetone before being baked overnight at 225°C. Teflon tubing was rinsed with hexane (Pesticide Grade, Fisher Scientific UK Ltd, Loughborough, UK) and allowed to air dry overnight.

To perform the bioassay, the Y-tube was placed horizontally on a solid bench, 1μl of an infected dog odour extract was injected onto the filter paper in one side of the apparatus and 1μl of an uninfected dog odour extract for comparison was injected into the other side of the apparatus. The holes in the wall of the Teflon tubes were sealed with PTFE ® tape

To test the response of sand-flies, each fly was removed from the holding cage and released individually at the open end of the stem of the Y-tube olfactometer. A timer was started and after three minutes the position of the sand fly within the Y-tube was recorded; either the test or control arm or, if it remained in the stem, a “no choice” was recorded. After 10 replicates, the rolled-up filter paper was removed and replaced, and to reduce any potential positional bias in the apparatus, the position of the test and control arms were swapped round by rotating the olfactometer through 180° in the horizontal plane. In total 80 female or 80 male sand flies were used per experimental replicate (i.e. each pair of infected and uninfected dog odour combinations were tested with 80 sand flies).

#### Sample selection

From the pool of available (n=133) dog hair odour samples, we selected the extracts collected from the hair of 15 infected and 15 uninfected dogs (n=15). Selection of the infected and uninfected dog hair odour extracts was based on PCR and qPCR analysis results. The selected infected dog hair extracts were from dogs with a parasite load that ranged from 1.3 ml^−1^ to 853.4 parasites ml^−1^ of blood. The uninfected dog hair extracts were from dogs with no detectable parasite DNA.

To determine if any attraction observed in the Y-tube experiments might be related to the discrimination value previously obtained from a VOC analyser (eNose) study [29] the 15 infected and 15 uninfected dog hair odour samples were also selected according to their discrimination values. Therefore, each infected and uninfected dog hair sample group had 5 high, 5 medium, 5 low discriminated individuals, and a cross comparison performed. The pairings are summarised in S1 Table.

### Study design

To carry out the study, the response of female (or male sand flies) to each infected dog hair odour sample was compared with their response to an uninfected dog hair sample in the Y-tube olfactometer. High, medium and low discriminated infected samples were cross compared with either a high, medium or low discriminated uninfected sample. The same 15 pairs of dog hair odour were used in both the female and male bioassays, although in a different order (S1 Table and S2 Table).

To avoid experimenter bias, the 15 infected and uninfected dog hair pairs were divided into 2 sub-groups. The infection status was known to the experimenter (unblinded) in one subgroup of nine dogs but unknown (blinded) in the other subgroup of 6 dogs.

All dog odour extracts were diluted by a factor of 1:10 to achieve optimal sand fly response [64]. Control bioassays were performed to determine female and male response to uninfected dog hair odour extract compared to a hexane control.

### Data Analysis

Binomial tests [65] were used to determine whether a greater number of female or male sand flies were attracted to infected dog odour extract than uninfected dog odour extract than would be expected by chance (50/50) for each experimental replicate. A binomial test was also used to compare response of female and male *Lu. longipalpis* to uninfected dog odour.

Response of female or male sand flies to infected or uninfected dog odour in the unblinded and blinded experiments was compared using two-tailed T-tests.

Response of female and male sand flies to the infected dog odour or uninfected dog odour was compared using two-tailed paired T-tests.

Response of female sand flies to infected dog odour extract was compared to response of male sand flies to infected dog odour extract using a two-tailed T-test.

Normality tests (Anderson Darling and Kolmogorov-Smirnov) were used to confirm that data were normally distributed and control data were analysed by binomial test.

To determine if there was a relationship between parasite load and the female and male sand fly response, we carried out regression analyses on the proportion of females (and males) that were attracted to the infected dog hair odour and the parasite load of the dog. A correlation analysis was used to determine if there was a relationship between the discrimination status of odour extracts obtained in the VOC analyser study and the proportion of female or male sand flies that responded to the extracts.

All statistical analyses were performed using MINITAB 19 (Minitab, Inc. 2020).

### Ethics

Dog blood and hair samples were taken from dogs that were also microchipped with the informed consent of their owners. Ethical approval was obtained from the Comissão de Ética no Uso de Animais (CEUA), Instituto Oswaldo Cruz (licence L-027/2017) in Brazil and Lancaster University Animal Welfare and Ethics Review Board (AWERB) in the UK. The CEUA approval complies with the provisions of Brazilian Law 11794/08, which provides for the scientific use of animals, including the principles of Brazilian Society of Science in Laboratory Animals (SBCAL). The AWERB approval complies with the UK Home Office guidelines of the Animals in Science Regulation Unit (ASRU) and in compliance with the Animals (Scientific Procedures) Act (ASPA) 1986 (amended 2012) regulations and was consistent with UK Animal Welfare Act 2006.

## Acknowledgments

We are grateful to Centro de Controle de Zoonoses in GV for permission to carry out the work as well as for their practical support. Ms M. Bell and Dr. C.F. de Souza, Lancaster University and Fiotec for providing technical support in GV. Dr R. Dillon for access to the *Lu. longipalpis* Jacobina colony at Lancaster University. Dr R.V. de Amaral for help with sand fly colony maintenance. We are profoundly grateful to the dog owners of Altinópolos, GV for allowing us to carry out this study on their cherished companion animals.

## Author roles

|  |  |
| --- | --- |
| Conceptualization | JGCH |
| Formal Analysis | MS, JGCH |
| Funding Acquisition | JGCH |
| Investigation | MS |
| Methodology | MS, JGCH |
| Project Administration | JGCH |
| Supervision | JGCH |
| Visualization | MS, JGCH |
| Writing – Original Draft Preparation | MS, JGCH |
| Writing – Review & Editing | MS, JGCH |

## Supporting information captions

S1 Table. **Female *Lutzomyia longipalpis* response to entrained odour of infected and uninfected dogs in unblinded and blinded Y-tube olfactometer experiments**. Summary of blinded and unblinded Y-tube bioassay experiments for female *Lu. longipalpis*. The odour extracts from infected dogs (in bold) and uninfected dogs are shown. Infected = the number of male sand flies attracted to the arm of the Y-tube containing infected odour; uninfected = the number of male sand flies attracted to the arm of the Y-tube containing uninfected odour; no-response = number of sand flies that did not respond in the Y-tube experiment. a = unblinded experimental replicates, b = blinded experimental replicates. Statistical significance was analysed by binomial test comparison of number of sand flies responding to test and control odours (\**P*≤0.05, \*\**P*≤0.01, \*\*\**P*≤0.001, \*\*\*\**P*≤0.0001).

S2 Table. **Male *Lutzomyia longipalpis* response to entrained odour of infected and uninfected dogs in the unblinded and blinded Y-tube olfactometer experiments.** Summary of blinded and unblinded Y-tube bioassay experiments for male *Lu. longipalpis*. The odour extracts from infected dogs (in bold) and uninfected dogs are shown. Infected = the number of male sand flies attracted to the arm of the Y-tube containing infected odour; uninfected = the number of male sand flies attracted to the arm of the Y-tube containing uninfected odour; no-response = number of sand flies that did not respond in the Y-tube experiment. a = unblinded experimental replicates, b = blinded experimental replicates. Statistical significance was analysed by binomial test comparison of number of sand flies responding to test and control odours (*ns* = not significant). The experimental outcomes are presented in the same order as for the female bioassays (Table 1) for ease of comparison.

## References

1. Maroli M, Feliciangeli MD, Bichaud L, Charrel RN, Gradoni L. Phlebotominae sandflies and the spreading of leishmaniases and other diseases of public health concern. Medical and Veterinary Entomology. 2013;27:25.

2. Burza S, Croft SL, Boelaert M. Leishmaniasis. The Lancet. 2018;392:20. doi: 10.1016/S0140-6736(18)31204-2.

3. Alvar J, Vélez ID, Bern C, Herrero M, Desjeux P, Cano J, et al. Leishmaniasis worldwide and global estimates of its incidence. PloS one. 2012;7(5):e35671–e. doi: 10.1371/journal.pone.0035671. PubMed PMID: 22693548.

4. WHO. Leishmaniasis Geneva: World Health Organisation; 2020 [updated 2 March 202005 May 2020]. Available from: https://www.who.int/en/news-room/fact-sheets/detail/leishmaniasis.

5. PAHO/WHO. Leishmaniasis. Epidemiological Report of the Americas, February 2018. 2018.

6. PAHO/WHO. Plan of action to strengthen the surveillance and control of Leishmaniasis in the Americas. 2017-2022. Department CDaHA; 2017.

7. Bezerra JMT, de Araújo VEM, Barbosa DS, Martins-Melo FR, Werneck GL, Carneiro M. Burden of leishmaniasis in Brazil and federated units, 1990-2016: Findings from Global Burden of Disease Study 2016. PLoS neglected tropical diseases. 2018;12(9):e0006697. doi: 10.1371/journal.pntd.0006697.

8. Courtenay O, Carson C, Calvo-Bado L, Garcez LM, Quinnell RJ. Heterogeneities in Leishmania infantum infection: using skin parasite burdens to identify highly infectious dogs. PLoS neglected tropical diseases. 2014;8(1):e2583. Epub 2014/01/15. doi: 10.1371/journal.pntd.0002583. PubMed PMID: 24416460; PubMed Central PMCID: PMCPMC3886905.

9. Brandão-Filho SP, Balbino VQ, Marcondes CB, Brazil RP, Hamilton JG, Shaw JJ. Should reproductively isolated populations of Lutzomyia longipalpis sensu lato receive taxonomically valid names? Memórias do Instituto Oswaldo Cruz. 2009;104:1197–200.

10. Solano-Gallego L, Miró G, Koutinas A, Cardoso L, Pennisi MG, Ferrer L, et al. LeishVet guidelines for the practical management of canine leishmaniosis. Parasites & Vectors. 2011;4(1):86. doi: 10.1186/1756-3305-4-86.

11. Lehane MJ. The Biology of Blood-Sucking in Insects. 2nd ed. Cambridge, UK: Cambridge University Press; 2005 2005.

12. Hamilton JGC, Hurd H. Parasite manipulation of vector behaviour. In: Lewis EE, Campbell, J.F., Sukhdeo, M.V.K., editor. The Behavioual Ecology of Parasites: CAB International; 2002. p. 384.

13. Hurd H. Manipulation of medically important insect vectors by their parasites. Annu Rev Entomol. 2003;48:141–61. Epub 2002/11/05. doi: 10.1146/annurev.ento.48.091801.112722. PubMed PMID: 12414739.

14. Gandon S. Evolution and Manipulation of Vector Host Choice. The American Naturalist. 2018;192(1):23–34. doi: 10.1086/697575. PubMed PMID: 29897804.

15. Mauck KE, De Moraes CM, Mescher MC. Effects of pathogens on sensory-mediated interactions between plants and insect vectors. Current Opinion in Plant Biology. 2016;32:53–61. doi: https://doi.org/10.1016/j.pbi.2016.06.012.

16. Koella JC, Sørensen FL, Anderson RA. The malaria parasite, Plasmodium falciparum, increases the frequency of multiple feeding of its mosquito vector, Anopheles gambiae. Proceedings Biological sciences. 1998;265(1398):763–8. doi: 10.1098/rspb.1998.0358. PubMed PMID: 9628035.

17. Lacroix R, Mukabana WR, Gouagna LC, Koella JC. Malaria infection increases attractiveness of humans to mosquitoes. PLoS Biol. 2005;3(9):e298. Epub 2005/08/04. doi: 10.1371/journal.pbio.0030298. PubMed PMID: 16076240; PubMed Central PMCID: PMCPMC1182690.

18. Batista EPA, Costa EFM, Silva AA. Anopheles darlingi (Diptera: Culicidae) displays increased attractiveness to infected individuals with Plasmodium vivax gametocytes. Parasites & Vectors. 2014;7(1):251. doi: 10.1186/1756-3305-7-251.

19. Cornet S, Nicot A, Rivero A, Gandon S. Both infected and uninfected mosquitoes are attracted toward malaria infected birds. Malaria journal. 2013;12:179-. doi: 10.1186/1475-2875-12-179. PubMed PMID: 23731595.

20. Díez-Fernández A, Martínez-de la Puente J, Gangoso L, López P, Soriguer R, Martín J, et al. Mosquitoes are attracted by the odour of Plasmodium-infected birds. International Journal for Parasitology. 2020. doi: https://doi.org/10.1016/j.ijpara.2020.03.013.

21. Busula AO, Bousema T, Mweresa CK, Masiga D, Logan JG, Sauerwein RW, et al. Gametocytemia and Attractiveness of Plasmodium falciparum-Infected Kenyan Children to Anopheles gambiae Mosquitoes. J Infect Dis. 2017;216(3):291–5. Epub 2017/09/02. doi: 10.1093/infdis/jix214. PubMed PMID: 28859429.

22. Cornet S, Nicot A, Rivero A, Gandon S. Evolution of plastic transmission strategies in avian malaria. PLoS Pathog. 2014;10(9):e1004308–e. doi: 10.1371/journal.ppat.1004308. PubMed PMID: 25210974.

23. Emami SN, Lindberg BG, Hua S, Hill SR, Mozuraitis R, Lehmann P, et al. A key malaria metabolite modulates vector blood seeking, feeding, and susceptibility to infection. Science. 2017;355(6329):1076–80. doi: 10.1126/science.aah4563.

24. O’Shea B, Rebollar-Tellez E, Ward RD, Hamilton JGC, El Naiem D, Polwart A. Enhanced sandfly attraction to Leishmania-infected hosts. Transactions of the Royal Society of Tropical Medicine and Hygiene. 2002;96(2):117–8. doi: 10.1016/s0035-9203(02)90273-7.

25. Nevatte TM, Ward RD, Sedda L, Hamilton JGC. After infection with Leishmania infantum, Golden Hamsters (Mesocricetus auratus) become more attractive to female sand flies (Lutzomyia longipalpis). Sci Rep. 2017;7(1):6104. Epub 2017/07/25. doi: 10.1038/s41598-017-06313-w. PubMed PMID: 28733676; PubMed Central PMCID: PMCPMC5522394.

26. De Oliveira LS, Rodrigues FM, de Oliveira FS, Mesquita PRR, Leal DC, Alcântara AC, et al. Headspace solid phase microextraction/gas chromatography–mass spectrometry combined to chemometric analysis for volatile organic compounds determination in canine hair: A new tool to detect dog contamination by visceral leishmaniasis. Journal of Chromatography B. 2008;875(2):6.

27. Magalhaes-Junior JT, Mesquita PR, Oliveira WF, Oliveira FS, Franke CR, Rodrigues Fde M, et al. Identification of biomarkers in the hair of dogs: new diagnostic possibilities in the study and control of visceral leishmaniasis. Anal Bioanal Chem. 2014;406(26):6691–700. Epub 2014/08/31. doi: 10.1007/s00216-014-8103-2. PubMed PMID: 25171830.

28. Magalhães-Junior JT, Oliva-Filho ADA, Novais HO, Mesquita PRR, M. Rodrigues F, Pinto MC, et al. Attraction of the sandfly Lutzomyia longipalpis to possible biomarker compounds from dogs infected with Leishmania infantum. Medical and Veterinary Entomology. 2019;33(2):322–5. doi: 10.1111/mve.12357.

29. Staniek ME, Sedda L, Gibson TD, de Souza CF, Costa EM, Dillon RJ, et al. eNose analysis of volatile chemicals from dogs naturally infected with Leishmania infantum in Brazil. PLoS neglected tropical diseases. 2019;13(8):e0007599. doi: 10.1371/journal.pntd.0007599.

30. Quinnell RJ, Courtenay O. Transmission, reservoir hosts and control of zoonotic visceral leishmaniasis. Parasitology. 2009;136(14):1915–34. doi: 10.1017/s0031182009991156. PubMed PMID: WOS:000273515200006.

31. Morrison AC, Ferro C, Pardo R, Torres M, Wilson ML, Tesh RB. Nocturnal activity patterns of Lutzomyia longipalpis (Diptera, Psychodidae) at an endemic focus of visceral leishmaniasis in Colombia. Journal of Medical Entomology. 1995;32(5):605–17. doi: 10.1093/jmedent/32.5.605. PubMed PMID: WOS:A1995RT60800007.

32. Spiegel CN, Dias DB, Araki AS, Hamilton JG, Brazil RP, Jones TM. The Lutzomyia longipalpis complex: a brief natural history of aggregation-sex pheromone communication. Parasit Vectors. 2016;9(1):580. Epub 2016/11/16. doi: 10.1186/s13071-016-1866-x. PubMed PMID: 27842601; PubMed Central PMCID: PMCPMC5109651.

33. Kelly DW, Dye C. Pheromones, kairomones and the aggregation dynamics of the sandfly Lutzomyia longipalpis. Animal Behaviour. 1997;53:721–31. doi: 10.1006/anbe.1996.0309. PubMed PMID: WOS:A1997WW45300005.

34. Bray DP, Hamilton JGC. Host odor synergizes attraction of virgin female Lutzomyia longipalpis (Diptera: Psychodidae). Journal of Medical Entomology. 2007;44(5):779–87. doi: 10.1603/0022-2585(2007)44[779:hosaov]2.0.co;2.

35. Jones TM, Hamilton JGC. A role for pheromones in mate choice in a lekking sandfly. Anim Behav. 1998;56(4):891–8. Epub 1998/12/16. PubMed PMID: 9790700.

36. Jones TM, Quinnell RJ. Testing predictions for the evolution of lekking in the sandfly, Lutzomyia longipalpis. Animal Behaviour. 2002;63:605–12. doi: 10.1006/anbe.2001.1946. PubMed PMID: WOS:000175391400022.

37. Borja LS, Sousa OMF, Solcà MDS, Bastos LA, Bordoni M, Magalhães JT, et al. Parasite load in the blood and skin of dogs naturally infected by Leishmania infantum is correlated with their capacity to infect sand fly vectors. Vet Parasitol. 2016;229:110–7. Epub 2016/11/05. doi: 10.1016/j.vetpar.2016.10.004. PubMed PMID: 27809965.

38. Borja LS, de Sousa OMF, Solca MD, Bastos LA, Bordoni M, Magalhaes JT, et al. Parasite load in the blood and skin of dogs naturally infected by Leishmania infanturn is correlated with their capacity to infect sand fly vectors. Veterinary Parasitology. 2016;229:110–7. doi: 10.1016/j.vetpar.2016.10.004. PubMed PMID: WOS:000388549200019.

39. Doehl JSP, Bright Z, Dey S, Davies H, Magson J, Brown N, et al. Skin parasite landscape determines host infectiousness in visceral leishmaniasis. Nature Communications. 2017;8. doi: 10.1038/s41467-017-00103-8. PubMed PMID: WOS:000404778800001.

40. Bray DP, Carter V, Alves GB, Brazil RP, Bandi KK, Hamilton JG. Synthetic sex pheromone in a long-lasting lure attracts the visceral leishmaniasis vector, Lutzomyia longipalpis, for up to 12 weeks in Brazil. PLoS neglected tropical diseases. 2014;8(3):e2723. Epub 2014/03/22. doi: 10.1371/journal.pntd.0002723. PubMed PMID: 24651528; PubMed Central PMCID: PMCPMC3961206.

41. Bell MJ, Sedda L, Gonzalez MA, de Souza CF, Dilger E, Brazil RP, et al. Attraction of Lutzomyia longipalpis to synthetic sex-aggregation pheromone: Effect of release rate and proximity of adjacent pheromone sources. PLoS neglected tropical diseases. 2018;12(12):e0007007. Epub 2018/12/20. doi: 10.1371/journal.pntd.0007007. PubMed PMID: 30566503; PubMed Central PMCID: PMCPMC6300254.

42. Matowo NS, Moore J, Mapua S, Madumla EP, Moshi IR, Kaindoa EW, et al. Using a new odour-baited device to explore options for luring and killing outdoor-biting malaria vectors: a report on design and field evaluation of the Mosquito Landing Box. Parasite Vector. 2013;6(1):137. doi: 10.1186/1756-3305-6-137.

43. Njiru BN, Mukabana WR, Takken W, Knols BGJ. Trapping of the malaria vector Anopheles gambiae with odour-baited MM-X traps in semi-field conditions in western Kenya. Malaria J. 2006;5(1):39. doi: 10.1186/1475-2875-5-39.

44. Jawara M, Smallegange RC, Jeffries D, Nwakanma DC, Awolola TS, Knols BGJ, et al. Optimizing Odor-Baited Trap Methods for Collecting Mosquitoes during the Malaria Season in The Gambia. Plos One. 2009;4(12):e8167. doi: 10.1371/journal.pone.0008167.

45. Rayaisse JB, Tirados I, Kaba D, Dewhirst SY, Logan JG, Diarrassouba A, et al. Prospects for the Development of Odour Baits to Control the Tsetse Flies Glossina tachinoides and G. palpalis s.l. Plos Neglect Trop D. 2010;4(3):e632. doi: 10.1371/journal.pntd.0000632.

46. WHO. Hamdbook for Integrated Vector Management. 2012.

47. Courtenay O, Dilger E, Calvo-Bado LA, Kravar-Garde L, Carter V, Bell MJ, et al. Sand fly synthetic sex-aggregation pheromone co-located with insecticide reduces the incidence of infection in the canine reservoir of visceral leishmaniasis: A stratified cluster randomised trial. PLoS neglected tropical diseases. 2019;13(10):e0007767. doi: 10.1371/journal.pntd.0007767.

48. Peel MC, Finlayson BL, McMahon TA. Updated world map of the Köppen-Geiger climate classification. Hydrol Earth Syst Sci. 2007;11(5):1633–44. doi: 10.5194/hess-11-1633-2007.

49. Climate-data.org. Climate data for cities worldwide – Climate: Governador Valadares 2017 [cited 2017 13th October 2017]. Available from: https://en.climate-data.org/info/sources/.

50. Barata RA, Peixoto JC, Tanure A, Gomes ME, Apolinario EC, Bodevan EC, et al. Epidemiology of visceral leishmaniasis in a reemerging focus of intense transmission in Minas Gerais State, Brazil. Biomed Res Int. 2013;2013:405083. Epub 2013/09/04. doi: 10.1155/2013/405083. PubMed PMID: 24000322; PubMed Central PMCID: PMCPMC3755404.

51. Valdivia HO, Almeida LV, Roatt BM, Reis-Cunha JL, Pereira AA, Gontijo C, et al. Comparative genomics of canine-isolated Leishmania (Leishmania) amazonensis from an endemic focus of visceral leishmaniasis in Governador Valadares, southeastern Brazil. Sci Rep. 2017;7:40804. Epub 2017/01/17. doi: 10.1038/srep40804. PubMed PMID: 28091623; PubMed Central PMCID: PMCPMC5238499.

52. Alves WA, Fonseca DS. Leishmaniose visceral humana: estudo do perfil clínico-epidemiológico na região leste de Minas Gerais, Brasil. J Health Biol Sci2018. p. 133–9.

53. Barata RA, Peixoto JC, Tanure A, Gomes ME, Apolinario EC, Bodevan EC, et al. Epidemiology of Visceral Leishmaniasis in a Reemerging Focus of Intense Transmission in Minas Gerais State, Brazil. Biomed Research International. 2013. doi: 10.1155/2013/405083. PubMed PMID: WOS:000323506000001.

54. SES-MG. Secretaria de Estado de Saúde de Minas Gerais. Programa de Vigilância e Controle da Leishmaniose Visceral. Boletim Epidemiológico, Leishmaniose Visceral Humana, Minas Gerais, 2010-2015. 2017. 8 p. 2017. Available from: http://vigilancia.saude.mg.gov.br/index.php/download/boletim-epidemiologico-leishmaniose-visceral-humana-minas-gerais-2010-2015/.

55. Brasil/SVS. Ministério da Saúde. Banco de dados do Sistema Único de Saúde - DATASUS, Informações de Saúde, Epidemiológicas e morbidade, Doenças e Agravos de Notificação (SINAN) 2020 [19/08/2020]. Available from: http://www2.datasus.gov.br/DATASUS/index.php?area=0203&id=29892192&VObj=http://tabnet.datasus.gov.br/cgi/deftohtm.exe?sinannet/cnv.

56. Pinheiro AD, da Costa ASV, de Oliveira RS, Reis MLC. Epidemiological aspects and spatial distribution of visceral leishmaniasis in Governador Valadares, Brazil, between 2008 and 2012. Revista Da Sociedade Brasileira De Medicina Tropical. 2020;53. doi: 10.1590/0037-8682-0216-2019. PubMed PMID: WOS:000504021800001.

57. Mary C, Faraut F, Lascombe L, Dumon H. Quantification of Leishmania infantum DNA by a real-time PCR assay with high sensitivity. J Clin Microbiol. 2004;42(11):5249–55. Epub 2004/11/06. doi: 10.1128/JCM.42.11.5249-5255.2004. PubMed PMID: 15528722; PubMed Central PMCID: PMCPMC525214.

58. Akella SV, Kirk WD, Lu YB, Murai T, Walters KF, Hamilton JG. Identification of the aggregation pheromone of the melon thrips, Thrips palmi. PLoS One. 2014;9(8):e103315. Epub 2014/08/08. doi: 10.1371/journal.pone.0103315. PubMed PMID: 25101871; PubMed Central PMCID: PMCPMC4125133.

59. Lawyer P, Killick-Kendrick M, Rowland T, Rowton E, Volf P. Laboratory colonization and mass rearing of phlebotomine sand flies (Diptera, Psychodidae). Parasite. 2017;24:42. Epub 2017/11/16. doi: 10.1051/parasite/2017041. PubMed PMID: 29139377; PubMed Central PMCID: PMCPMC5687099.

60. Moraes CS, Aguiar-Martins K, Costa SG, Bates PA, Dillon RJ, Genta FA. Second Blood Meal by Female Lutzomyia longipalpis: Enhancement by Oviposition and Its Effects on Digestion, Longevity, and Leishmania Infection. Biomed Res Int. 2018;2018:2472508. Epub 2018/05/18. doi: 10.1155/2018/2472508. PubMed PMID: 29770328; PubMed Central PMCID: PMCPMC5889884.

61. Ward RD, Lainson R, Shaw JJ. Some methods for membrane feeding of laboratory reared, neotropical sandflies (Diptera: Psychodidae). Ann Trop Med Parasitol. 1978;72(3):269–76. Epub 1978/06/01. doi: 10.1080/00034983.1978.11719315. PubMed PMID: 666397.

62. Denlinger DS, Li AY, Durham SL, Lawyer PG, Anderson JL, Bernhardt SA. Comparison of In Vivo and In Vitro Methods for Blood Feeding of Phlebotomus papatasi (Diptera: Psychodidae) in the Laboratory. J Med Entomol. 2016;53(5):1112–6. Epub 2016/06/15. doi: 10.1093/jme/tjw074. PubMed PMID: 27297215.

63. Meireles-Filho ACA, Kyriacou CP. Circadian rhythms in insect disease vectors. Memorias do Instituto Oswaldo Cruz. 2013;108 Suppl 1(Suppl 1):48–58. doi: 10.1590/0074-0276130438. PubMed PMID: 24473802.

64. Nevatte TM. Changes in Golden Hamster behaviour and attractiveness to the sand fly Lutzomyia longipalpis as a result of Leishmania infection. UK: Keele University; 2006.

65. Sokal RR, Rohlf FJ. Biometry: the principles and practice of statistics in biological research. 3rd ed. New York: W.H. Freeman and Company; 1995.

