## Supplemental Table 1 for "Odour of domestic dogs infected with *Leishmania infantum* is attractive to female but not male sand flies: evidence for parasite manipulation"

| **expt** | **Dog pair** | **infected odour** | **un-infected odour** | **no-response** | ***P*** |
| --- | --- | --- | --- | --- | --- |
| **1** | **176** vs 021^a^ | 49 | 21 | 10 | *** |
| **2** | **141** vs 137^a^ | 46 | 21 | 13 | *** |
| **3** | **178** vs 181^a^ | 51 | 18 | 11 | **** |
| **4 stuck at the top of page4** | **105** vs 037^a^ | 44 | 24 | 12 | ** |
| **5** | **140** vs 004^a^ | 51 | 20 | 9 | **** |
| **6** | **003** vs 130^a^ | 42 | 23 | 15 | ** |
| **7** | **074** vs 093^a^ | 40 | 24 | 16 | * |
| **8** | **082** vs 175^a^ | 45 | 26 | 9 | ** |
| **9** | **102** vs 124^a^ | 42 | 28 | 10 | * |
| **10** | **126** vs 169^b^ | 48 | 25 | 7 | ** |
| **11** | **047** vs 153^b^ | 45 | 26 | 9 | ** |
| **12** | **044** vs 005^b^ | 48 | 26 | 6 | ** |
| **13** | **080** vs 136^b^ | 45 | 25 | 10 | ** |
| **14** | **134** vs 070^b^ | 42 | 29 | 9 | * |
| **15** | **019** vs 043^b^ | 47 | 22 | 11 | *** |
|  | **sum** | **685** | **358** | **157** |  |
|  | **mean±se** | **45.7±0.9** | **23.9±0.8** | **10.5±0.7** |  |
