## Supplemental Table 2 for "Odour of domestic dogs infected with *Leishmania infantum* is attractive to female but not male sand flies: evidence for parasite manipulation"

| **expt** | **dog** | **Infected**  **odour** | **un-infected odour** | **no-response** | ***P*** |
| --- | --- | --- | --- | --- | --- |
| **1** | **176** vs 021^a^ | 35 | 37 | 8 | *ns* |
| **10** | **141** vs 137^b^ | 34 | 36 | 10 | *ns* |
| **11** | **178** vs 181^b^ | 36 | 36 | 8 | *ns* |
| **6** | **105** vs 037^a^ | 34 | 38 | 8 | *ns* |
| **8** | **140** vs 004^a^ | 37 | 37 | 6 | *ns* |
| **7** | **003** vs 130^a^ | 37 | 36 | 7 | *ns* |
| **5** | **074** vs 093^a^ | 36 | 34 | 10 | *ns* |
| **3** | **082** vs 175^a^ | 38 | 31 | 11 | *ns* |
| **15** | **102** vs 124^b^ | 39 | 32 | 9 | *ns* |
| **13** | **126** vs 169^b^ | 34 | 38 | 8 | *ns* |
| **14** | **047** vs 153^b^ | 35 | 38 | 7 | *ns* |
| **9** | **044** vs 005^a^ | 35 | 38 | 7 | *ns* |
| **2** | **080** vs 136^a^ | 35 | 35 | 10 | *ns* |
| **12** | **134** vs 070^b^ | 38 | 35 | 7 | *ns* |
| **4** | **019** vs 043^a^ | 38 | 35 | 7 | *ns* |
|  | **sum** | **541** | **536** | **123** |  |
|  | **mean±se** | **36.1±0.4** | **35.7±0.6** | **8.2±0.4** |  |
